# Interplay of immigration and coevolution constrains the structure of specialist-generalist communities

**DOI:** 10.1101/2025.08.27.670321

**Authors:** Vasco J. Lepori, Nicolas Loeuille, Rudolf P. Rohr

**Affiliations:** Department of Biology, University of Fribourg, CH-1700 Fribourg, Switzerland; Institute of Terrestrial Ecosystems, D-USYS, ETH Zürich, CH-8092 Zürich, Switzerland; Sorbonne Université, UPEC, CNRS, IRD, INRAE, iEES-Paris, F-75005 Paris, France

## Abstract

Species and populations in nature fall along a specialist-generalist continuum in habitat preferences and resource use. Despite a rich literature on the evolution of specialization at the population level, most studies focusing on community assembly, species packing and diversification have considered solely niche positions within a community. Yet, empirical evidence suggests that generalism levels in communities respond dynamically to the current biotic environment and species interactions. Here, we jointly consider niche positions and generalism, and ask how different modes of community assembly from evolutionary assembly to repeated immigration shape the resulting community. We find that coevolution can lead to diversification, including spontaneous emergence of asymmetric configurations of generalist and specialist phenotypes. We further observe distinct alternative evolutionary endpoints, comprising communities consisting of either many specialists or fewer generalists, some of which are only locally uninvasible. More broadly, we show how the history of local coevolution and dispersal events jointly influence community dynamics and structure. By modeling assembly along a continuum from pure coevolution to pure dispersal, we highlight how high rates of evolution constrain richness while promoting generalism, partly because evolutionary assembly can get stuck at local optima. In contrast, communities shaped by immigration tend to be more species-rich, with diverse and specialist characters. Finally, higher rates of evolution decrease invasion success in mixed-assembly scenarios, a pattern consistent with community monopolization. We discuss how our theoretical findings match observational and experimental evidence for community assembly across scales.

## Introduction

In nature, species and populations differ not only in their environmental optima, but also in the breadth and variability of the resources they consume and environmental conditions they tolerate. This concept is encapsulated within the classical notion of an ecological niche, which can be characterized by both an optimum and a width around said optimum. In empirical investigations of niche width, species and populations are observed to span a spectrum from specialists to generalists in natural ecosystems (Forister et al., 2015; Hulburt, 1985; Thomas et al., 2016). This variation affects their ecology, their ability to withstand environmental variability, and their likelihood of suffering from secondary extinctions (Brodie et al., 2014).

Theoretical and empirical studies have further underscored the dynamic nature of niche width, due to genetic change or phenotypic plasticity, which play a role in adaptation to environmental variation (Kassen, 2002). Generalism can, for example, help buffer temporal environmental variability (Ketola et al., 2013; Reboud and Bell, 1997), or act as a risk-spreading mechanism through diversification of resource use (portfolio effect or bet-hedging), although finding good empirical evidence for this effect is challenging (Childs et al., 2010). Evolutionary models of consumer generalism have often focused on consumer specialization on two discrete resources (e.g. Egas et al. (2004); Kisdi (2002); Rueffler et al. (2007); Wickman et al. (2019)) and stressed the importance of allocation trade-offs for population-level generalism. This idea is reflected in Levins’ fitness set concept, which states that the curvature of the fitness values across different environments or resources (i.e. the fitness cost in habitat 1 in order to perform well in habitat 2) determines whether populations evolve towards jack-of-all trades or highly specialized strategies (Levins, 1962).

This idea is echoed in the work of Ackermann and Doebeli (2004), which employed an interesting alternative approach to modeling the evolution of generalism: rather than treating resources as discrete categories, they represented consumer niche width along a continuous resource axis, adapting the classic models of resource-axis packing (Dieckmann and Doebeli, 1999; MacArthur, 1969). These classical theories, which assumed constant niche widths, suggest that the pool of resources for which species compete determines available niches and ultimately sets an upper bound on diversity (MacArthur, 1969). Empirical examples similarly show that niche filling limits diversification events in both natural (Price et al., 2014) and experimental systems (Brockhurst et al., 2007), and decreases the invasion success of newcomers (Eisenhauer et al., 2013). Niche widths also play a role in community dynamics: when species are more generalist, their interactions within the community are more widespread. The correlation between niche width and the number of realized interspecific interactions (potential and realized generalism *sensu* Robinson and Strauss (2020)) has been observed in mutualistic communities (Bascompte, 2013). Conversely, the level of specialization can adapt to the biotic environment: Parent et al. (2014) observed a reduction in niche width under increased competition in an experiment using flour beetles, while in Jones and Post (2013) increased competition lead to narrower niches in alewife populations (this effect was mitigated by enhanced resource availability). The effect of resources was confirmed by Robinson and Strauss (2020) who showed that generalist herbivores exhibit a narrower diet breadth in low-resource soils, and highlighted the direct consequences this had on community assembly and structure. Finally, Costa-Pereira et al. (2019) also documented how niche widths in frogs increase with resource availability, and contract when species diversity is higher. All these examples suggest complex interactions between resource availability, generalism, and diversity, shaping patterns of niche occupancy.

In an effort to model these patterns, Barabás and D’Andrea (2016) showed how high intraspecific variation negatively impacts community richness. However, as noted by Wickman et al. (2023), Quantitative Genetics approaches are limited in their ability to model community assembly because species richness is fixed at the start rather than being an outcome of the model (but see recent advances by Wickman et al. (2023) and Lion et al. (2023)). Hence, when both species richness and generalism vary during community assembly through evolutionary (e.g. diversification) and ecological processes (e.g. immigration, Vellend (2008)), the determinants of community structure remain an open question. Moreover, the interplay between local evolution and dispersal can greatly alter community assembly (De Meester et al., 2016; Leibold et al., 2022b). For instance, rapid adaptive evolution in *Daphnia magna* leads to cascading changes and shifts final zooplankton community composition compared to communities with non-adapted *Daphnia* genotypes (Pantel et al., 2015). Conversely, in a bacterial model system, diversification was shown to depend on the timing of invasion by a different strain (Fukami et al., 2007). Relatedly, rapid evolution of resident species to occupy free niches has been shown to prevent subsequent invasions by potential competitors. This effect, named community monopolization, has first been observed in natural communities (De Meester et al., 2002; Gillespie, 2004), subsequently formalized in a series of theoretical articles (Loeuille and Leibold, 2008; Urban and De Meester, 2009), and finally demonstrated in experimental settings (Farkas et al., 2013).

We here investigate eco-evolutionary assembly in generalist-specialist communities, to reveal how concurrent shifts in generalism and niche positions affect niche occupancy patterns under different assembly scenarios. We model coevolution of niche position and width as in Ackermann and Doebeli (2004), but in a species-rich context and with a focus on community assembly. Generalism and community diversity here both emerge as outcomes of the assembly process. We expect to observe diversification in sufficiently large resource landscapes (Ackermann and Doebeli, 2004; Doebeli and Dieckmann, 2000), however, the extent to which the evolution of generalism will offset this effect remains unclear (Barabás and D’Andrea, 2016). We assess how allocation trade-off and modes of community assembly act on this tension, by conducting simulations across a range of scenarios including purely evolutionary community assembly, successive immigration events, and mixed-assembly modes.

## Ecological dynamics: a Lotka-Volterra model for consumer competition

We consider a community of *S* consumer species that exploit a continuum of resource types along an axis *x*, with each resource type regenerating at a rate *ϕ*(*x*). Consumer feeding niches are characterized by a position *µ_i_* (their optimal resource type) and a width *σ_i_* (the spread of resources they can utilize efficiently). The consumption function of species *i* at any location on the resource axis is thus *f_i_*(*x*) = *f* (*x|µ_i_, σ_i_*), which we take to follow a Gaussian form. If we assume that resource dynamics are faster than consumer dynamics, we can describe the abundances of the consumer community using the well known Lotka-Volterra competition model shown in Equation 1 (Volterra, 1931). Consumer-consumer competition coefficients are given by their overlap in resource use across the axis *α_ij_* := ∫ *f_i_*(*x*)*f_j_*(*x*)*dx*. Consistent with the limiting similarity principle, competition becomes more intense the more similar the resources they exploit (Macarthur and Levins, 1967; Taper and Case, 1992). Consumer intrinsic growth rates *r_i_* depend on the continuous sum of resources acquired at each location *x* on the resource axis, minus a fixed mortality term *m* that ensures consumers whose feeding rates approach zero are deterministically removed: *r_i_* := *−m* + ∫ *ϕ*(*x*)*f_i_*(*x*)*dx*. We also assume that transformation of consumed resources into consumer biomass occurs with perfect efficiency.

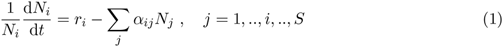

We focus our attention on the simple case of a bounded uniform resource regeneration function, with *ϕ*(*x*) = *h* between [*−l,* +*l*], and *ϕ*(*x*) = 0 everywhere else (boxcar function). Integrals for *α* are thus computed on the closed interval [*−l,* +*l*], so that consumers only compete for existing resources. We finally obtain the following model of consumer competition from equation (1), also shown graphically in Figure 1:

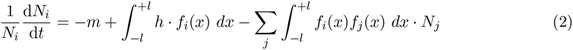

**Figure 1:**
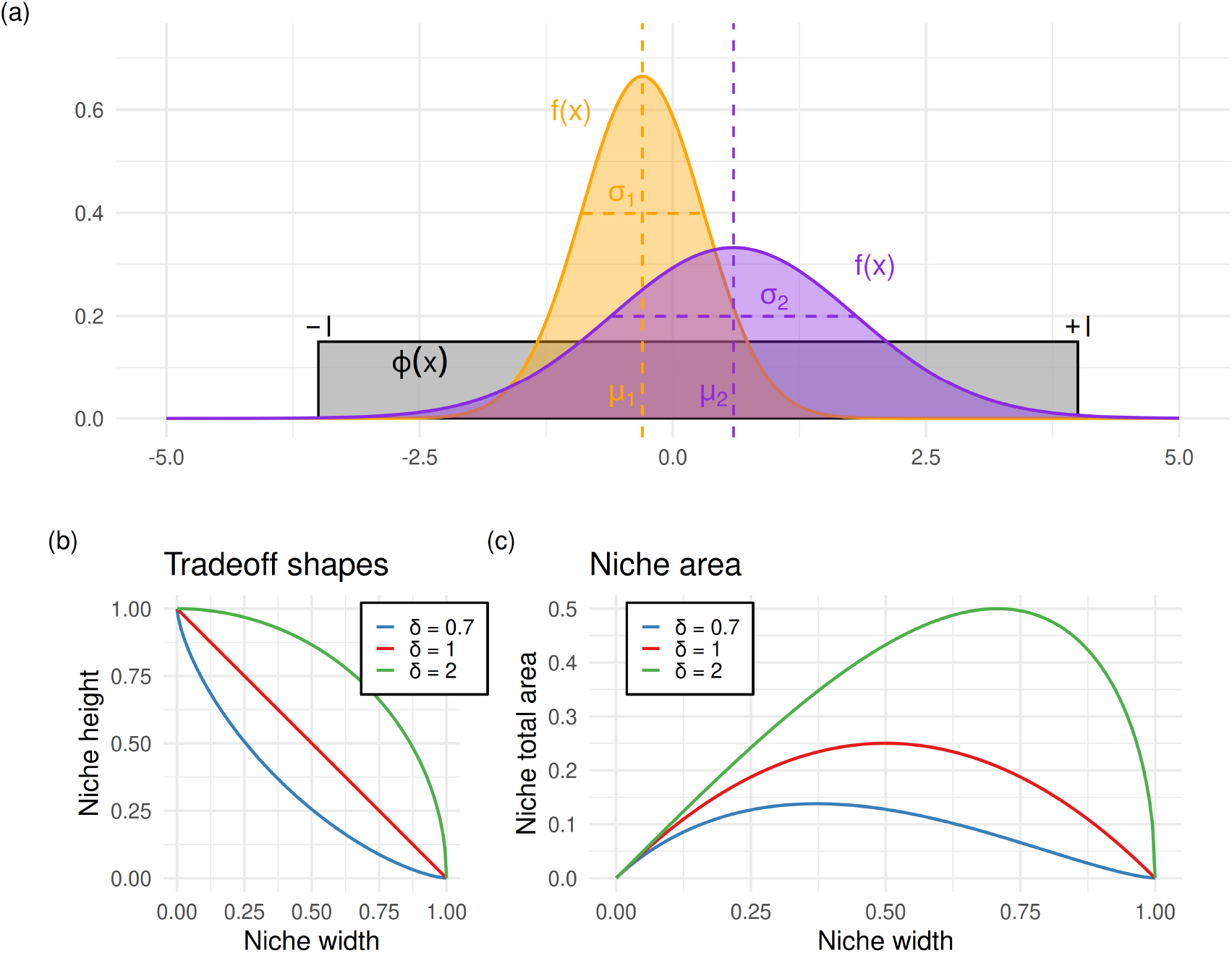
A competition model for generalist-specialist communities. (a) Graphical representation of consumer niches *f* (*x|µ, σ*) for a specialist (orange) and a generalist phenotype (purple) as defined by their niche position or optimum *µ* and their niche width *σ* on the resource axis. (b) trade-off shapes regulate the cost in niche amplitude to achieve a certain niche width: strong trade-offs (blue) impose a high cost to niche height to achieve a large niche width. (c) Note that this trade-off function does not preserve niche areas: for a given value *δ*, the optimal niche width value maximizing niche area is: argmax*_σ_ A_f_* = 2*^−^*^1^*^/δ^*. A summary of all model parameters is available in Table 1.

**Table 1.** Summary of symbols used.

| Symbol | Variable or parameter |
|---|---|
| *N<sub>i</sub>* | abundance of consumer *i* |
| *x* | a type of resource, location on resource axis |
| *f<sub>i</sub>*(*x*\|*μ<sub>i</sub>*, *σ<sub>i</sub>*) | feeding niche of consumer *i* |
| *μ<sub>i</sub>* | niche position of consumer *i*, optimal resource |
| *σ<sub>i</sub>* | niche width, generalism of *i* |
| *a<sub>i</sub>* | niche height of consumer *i* |
| *δ* | cost of generalism (trade-off strength) |
| *m* | intrinsic mortality rate of consumers |
| [−*l*, +*l*] | resource domain on the *x* axis |
| *h* | regeneration rate of resources within [−*l*, *l*] |
| *r<sub>i</sub>* | growth rate of consumer *i* |
| *α<sub>ij</sub>* | p.c. competition effect of *j* on *i* |
| *ω*(*μ*′, *σ*′) | relative fitness function of a mutant |

Appendix S1 details how this model can be derived from a bipartite resource-consumer model where resources are explicitly considered as state variables (original derivation by MacArthur 1972). It is worth noting that the Lotka-Volterra formulation is fully equivalent to the explicit model only in cases where no resources become extinct. When extinctions occur, the two formulations start to differ quantitatively; however, as we show in Appendix S2, our main findings remain qualitatively robust.

We model the specialism-generalism continuum as varying niche widths *σ* in the gaussian-shaped consumption niche *f_i_*(*x*) = *a_i_* exp[*−*(*x − µ_i_*)^2^*/*(2*σ_i_*)]. Here, we introduce a cost to being a generalist, to prevent the emergence of a Darwinian demon, a phenotype that could use any resource with perfect efficiency, out-competing all others. We materialize this cost as a trade-off between a consumer niche’s width *σ_i_* and its peak height *a_i_*, which we model on the [0, 1] interval. We select a flexible shape defined as *σ^δ^* + *a^δ^* = 1, which enables us to examine the importance of the assumed trade-off shape. At smaller values of *δ*, large niche widths come at the cost of performing poorly across most environments (Figure 1, panel B), whilst larger values of *δ* allow for greater generalism with lower performance costs in the preferred environment. We explore different trade-off forms because very little empirical information exists on the actual shape of these tradeoffs across clades, yet theoretical studies have revealed how their convexity can crucially determine evolutionary dynamics, and specifically the evolutionary stability of strategies (De Mazancourt and Dieckmann, 2004; Kisdi, 2006). We expect evolution of more generalist phenotypes for weak tradeoffs (concave, larger values of *δ*), and more diversification for stronger, convex trade-offs (smaller values of *δ*, Fig 1b)).

## Eco-evolutionary dynamics of consumer traits

To study eco-evolutionary dynamics of niche position and width in communities, we consider the per-capita growth rate when rare *ω_i_* of a mutant morph with traits 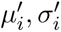 in a community of residents *j* = 1*, …, S*. We assume that mutations are rare compared to ecological dynamics, so that residents abundances reach their equilibrium 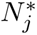 between mutation events. These assumptions yield the following fitness definition, which is inherently density- and frequency-dependent (Metz et al., 1992):

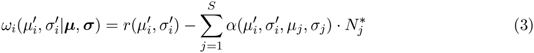

By further assuming very small mutational steps, we can study the coevolution of niche positions *µ_i_* and niche width *σ_i_* using the canonical equation of adaptive dynamics (Dieckmann and Law, 1996). The rates and direction of evolution over time (the fitness gradients) are given by the derivatives of mutant fitness with respect to each trait, multiplied by the population size *N* and a constant *M* encapsulating both the frequency and amplitude of mutations.

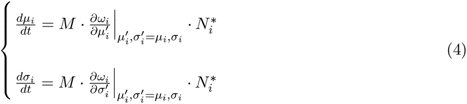

There is one equation per trait and per morph, and we assume the same evolutionary rates *M* for the two traits throughout the main text. This assumption appears not to influence evolutionary equilibria, which are determined solely by fitness gradients, but differing evolutionary rates between the two traits can slightly alter transient community dynamics (shown in Appendix S4). Furthermore, we assume that there are no physiological or genetic constraints that would cause evolutionary changes in the optimal resource type of species *i* to be inherently associated with changes in its generalism. This independence assumption is equivalent to saying that the covariance matrix in the coevolution equation is the identity matrix (Leimar, 2009; Metz, 2011).

A singular strategy *µ^∗^*, *σ^∗^*, or a coalition of strategies for polymorphic cases, corresponds to an evolutionary endpoint where directional evolution over time comes to a halt, i.e. the fitness gradients become null:

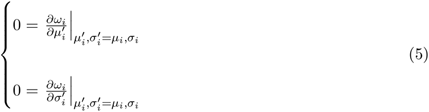

At such a location, the evolutionary strategy may either be a stable, uninvasible endpoint of evolution, or it could be a local fitness minimum where morphs experience disruptive selection and diversification can occur (Geritz et al., 1998). To evaluate evolutionary stability of the morph *i*, we compute the Hessian matrix of second derivatives as follows.

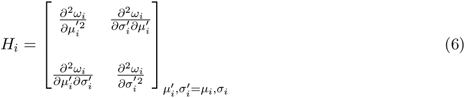

This matrix describes the curvature of the fitness function and must be negative-definite (both eigenvalues of *H_i_* must be negative) for the strategy to be an uninvasible evolutionary endpoint (ie, an ESS) in the morph *i* (Ackermann and Doebeli (2004); Leimar (2009)). Note that there is one hessian matrix per morph, and if all morphs are evolutionarily stable, the community itself is an Evolutionarily Stable Community (Edwards et al., 2018).

## Monomorphic evolutionary dynamics

We start by analyzing single-morph evolution (Figure 2a). A single consumer evolves towards a strategy that is always located in the center of the resource axis to efficiently access many resources (*µ^∗^* = 0, this is shown mathematically in Appendix S3), while its generalism level *σ^∗^* depends on the width of the resource axis and the trade-off strength. We already observe the emerging tension between generalism and diversification: increased resources lead to higher generalism evolution but eventually destabilize the strategy, ultimately leading to diversification (Figure 2b). At fixed resource availability, trade-off strength constrains the evolution of generalism. Lower costs (high *δ*) promote stable, generalist strategies, while strong trade-offs (smaller *δ*) lead to narrower niche widths, disruptive selection and diversification into multiple specialists.

**Figure 2:**
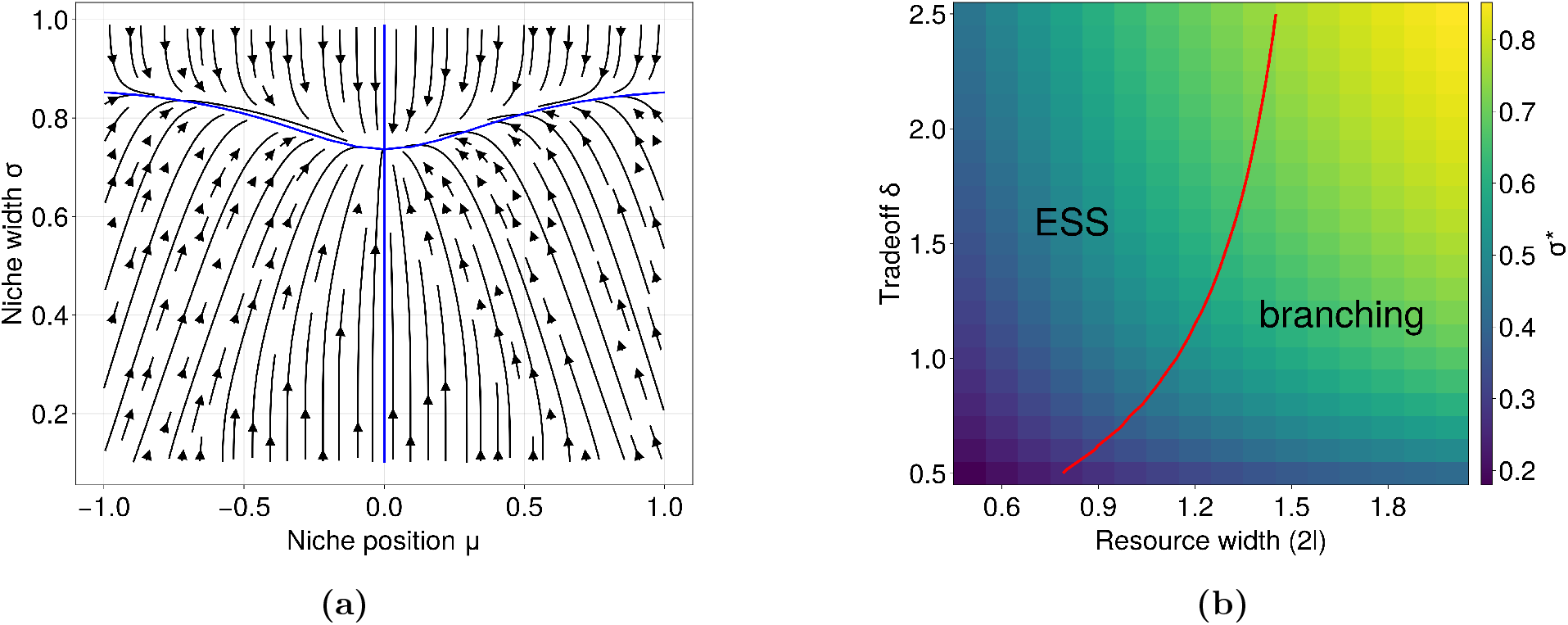
Evolutionary dynamics for a single-morph population. (a) Evolutionary trajectories converge to a singular strategy located in the center of the resource domain *µ^∗^* = 0. The blue lines indicate the isoclines where the evolutionary gradient becomes null for either niche position or width (Equ.5), which intersect at the singular strategy. Parameters for this trajectory are resource width 2*l* = 1.8*, δ* = 1.5. Panel (b) shows how both the value and stability of the generalism strategy *σ^∗^* vary with the resource scenario and trade-off strength. The red line determines the boundary of evolutionary stability, and is computed as the zero-contour line for the leading eigenvalue of the hessian matrix (Equ.6). Other parameters shared by both panels are *h* = 5*, m* = 0.2.

## Past the branching point: polymorphic dynamics and evolutionary endpoints

When resources are sufficiently diverse or the cost of generalism sufficiently high, we observe diversification at the singular strategy. We explore these polymorphic evolutionary dynamics by numerically solving equation (4) until a new end configuration is reached, upon which we re-evaluate evolutionary stability (invasibility). If unstable, we add a new morph at a random position nearby (in line with the small mutations assumption). We repeat this process until we reach a point that is uninvasible in the neighborhood of all resident morphs. For a given resource and trade-off scenario, we perform multiple simulations with randomized initial conditions. Initial values of *σ* and *µ* are drawn from uniform distributions and, to promote sampling and discovery of high-richness evolutionary endpoints, half the replicate runs are initialized at highly rich randomized feasible community, while in the other half the initial diversity is also drawn uniformly. All numerical simulations were carried out in Julia v1.12, and outputs were plotted using R v4.6.

This procedure reveals interesting dynamics, including the emergence of generalist-specialist coalitions, and alternative stable endpoints of evolution (Figure 3). Due to randomness in initial conditions and branching events, the evolutionary process may give rise to different communities, as illustrated in Figure 3, where either a dimorphic coalition of strategies (specialist-generalist, panels A,C) or a triplet (two specialists and one generalist, panels B,D) is obtained for the same parameter set. Importantly, while both of these configurations are locally evolutionarily stable (no further diversification ensuing), only the 3 morph solution is globally uninvasible. The two-morph coalition is, in fact, susceptible to invasion in certain regions of the fitness landscape that are unreachable by small mutations (in blue in Figure 3c) but could be accessible through immigration. Only the most species-rich of these attractors is a globally uninvasible polymorphic strategy (a strict Evolutionarily Stable Community *sensu* Edwards et al. (2018)), while the undersaturated end states are local but non-global ESCs where outside immigration could still lead to further community assembly.

**Figure 3:**
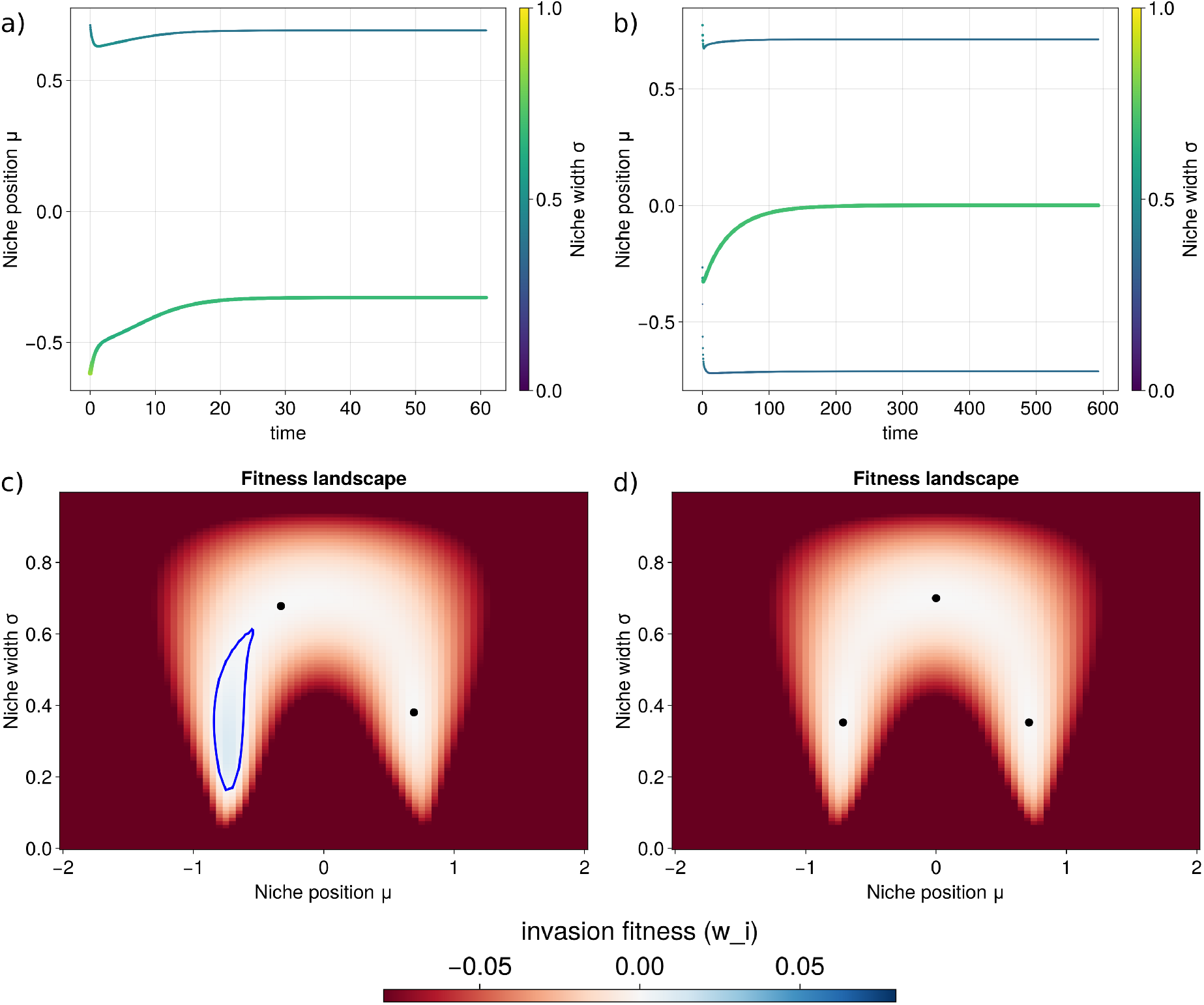
Two alternative endpoints of evolution for identical ecological conditions. Panels (a-b): the x-axis represents time, the y-axis shows niche positions, and color represents niche widths. The final invasion landscapes (c-d) picture mutant fitness as a function of mutant traits *µ* and *σ*. The 2-morph combination (a,c) is a non-global ESS which is only uninvasible in the neighborhood of the resident traits (in black in panel c), and the area of positive invasion fitness delimited by the blue line corresponds to trait combinations that would successfully invade the community. Conversely, the 3-morph coalition is globally uninvasible (panels b,d). The parameters for this pair of simulations are the same, and equal to *δ* = 2.0, resource width 2*l* = 1.7, *m* = 0.2, *h* = 5.

Figures 4 and 5 show a more comprehensive view of community organization patterns (species packing) in increasingly wide resource landscapes. While evolved richness increases with resource availability, the trade-off shape also affects community structure at higher levels of richness. When costs of generalism are higher, evolution favors diversification leading to richer, less generalist communities for a given resource landscape (Figure 4a). Moreover, while typical fixed-width models show a monotonic increase in community richness with larger resources, we here observe alternative evolutionary states, as depicted in Figure 5. These states lie on a trade-off between (average) generalism and richness (Figure 4c), with either many specialists or fewer generalist morphs. Of these communities, the richest ones are globally uninvasible Evolutionarily Stable Communities, while the invasible end-states are less species-rich, but tend to be more generalist.

**Figure 4:**
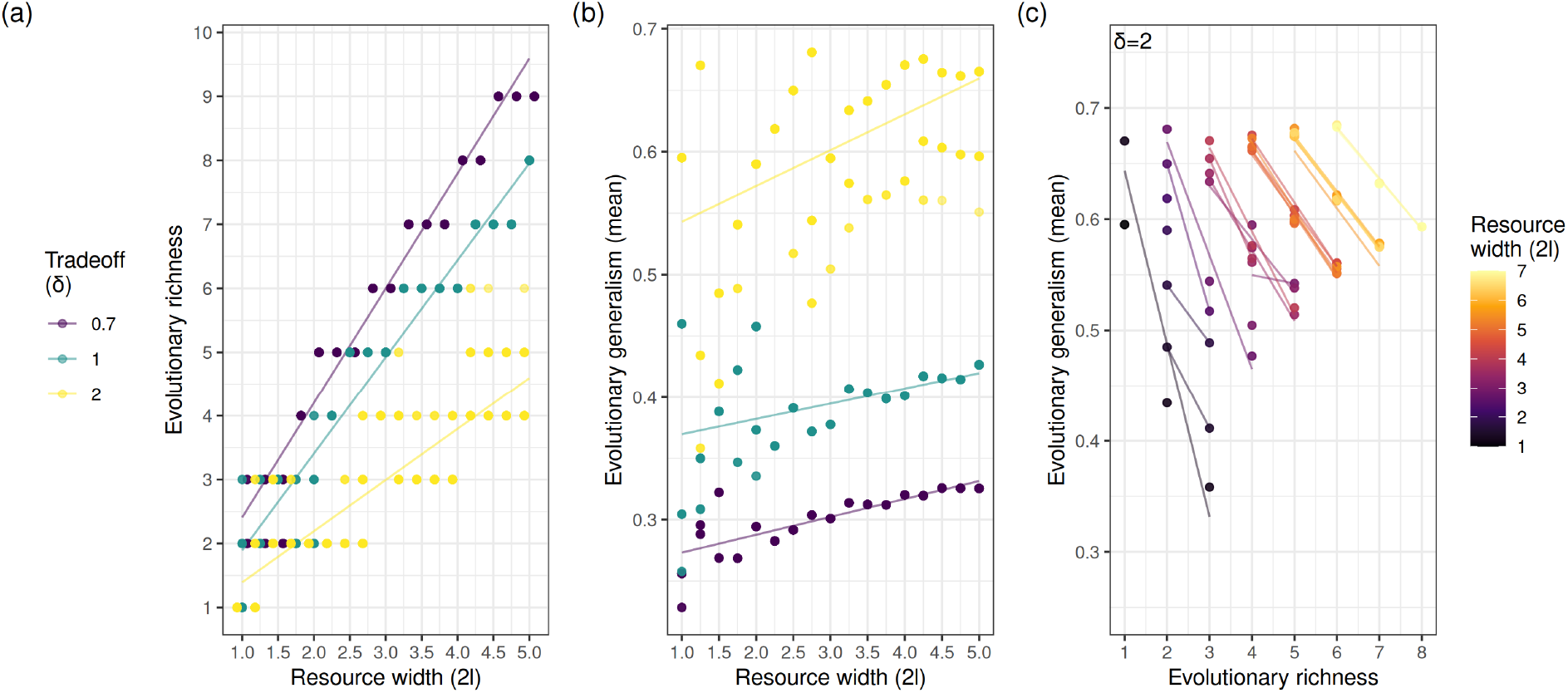
Evolutionary richness depends on resource availability and trade-off strength. Each dot is an an evolutionary endpoint reached from a randomly initialized community. Panel a) shows the increase in evolutionary richness with resource axis width for different trade-off strengths *δ*. Panel (b): stronger trade-offs also yield more specialized communities. Panel (c): richness and generalism of the alternative evolutionary stable communities shown in Figure 5, across resource levels and *δ* = 2. Lines are linear fits within each group. We ran 20 replicates for each combination of *δ* and *l*, and 50 replicates for each value of *l* in panel c). Shared parameters for all three panels: *h* = 5*, m* = 0.2

**Figure 5:**
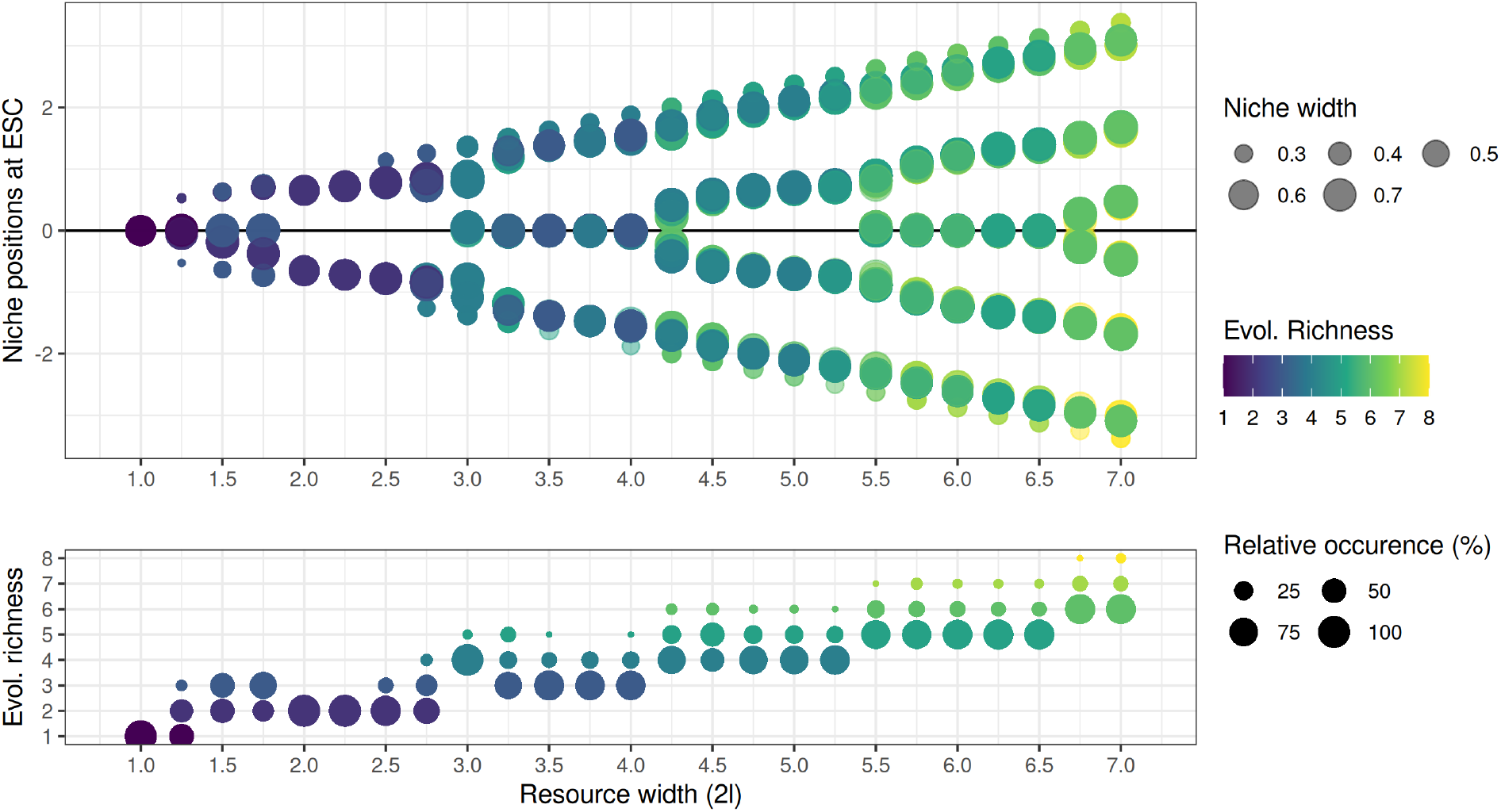
Alternative evolutionary endpoints and their frequency along a resource availability gradient. Top: all evolutionary communities found at each resource width *l*. We ran 100 replicates for each resource level, starting from randomized initial communities. Each dot is a morph, colors indicate the number of morphs in a same coalition (e.g. yellow corresponds to 8-morph evolutionary communities) and dots sizes indicate their niche width (*σ^∗^*). Alternative coalitions (or the same resource and trade-off levels may have the same richness but different arrangements (e.g., one symmetric and one asymmetric). Bottom: frequency of each configuration at each resource level (colors as above). Note that frequencies are solely illustrative and depend on the simulation method and initial conditions. Parameters: *δ* = 2.0*, h* = 5*, m* = 0.2.

## Assembly regimes influence community structure

The fact that evolution often leads to communities where further assembly would only be possible through immigration of novel traits underscores how different assembly modes can crucially affect community structure. To investigate this, we compared how communities assembled by pure evolution and diversification (small mutations and branching) differ from communities assembled by sequential invasions of migrants from a regional pool, and from intermediate communities shaped by both forces simultaneously. In the successive invasion scenario, arrivals are modeled as a Poisson process with rate *λ*, so that inter-arrival times are exponentially distributed. Newcomers are drawn from a regional pool modeled as samples from a 2-D Uniform distribution *µ ∼* Uniform(*−l,* +*l*) and *σ ∼* Uniform(0, 1). They can invade the resident community if their per capita growth rate is positive (1*/N · dN/dt >* 0), and can also make residents go extinct (we consider a consumer morph extinct when their abundance falls below *N <* 10*^−^*^4^). At the other end of the spectrum, we generate evolutionary assemblages by pure adaptive evolution and diversification (i.e. small mutations) from low initial richness, following the procedure described earlier. Along this gradient, we generate communities along the evolution-immigration spectrum by modulating both the rate of immigration *λ* (mean rate of arrivals per unit time) and the rate of evolution *M* (assumed identical for both traits, but see Appendix S4). Ecological dynamics are always at equilibrium between immigration events, reflecting the assumption that they are faster than both evolution and immigration (consecutive assembly scenario *sensu* Leibold et al. (2019)).

Figure 6 illustrates the effects of relative rates of evolution and immigration on community structure. Communities purely shaped by immigration (yellow in Figure 6) exhibit high richness and high levels of specialism, as well as fast turnover. These communities are all ecologically stable, but they are not endpoints of evolution: without a constant influx of immigrants, they would lose diversity over time through competitive exclusion caused by converging evolution Edwards et al. (2018)). Similarly, high-immigration, low-mutation communities can transiently show a pattern of oversaturation, but they eventually reach the globally uninvasible state (Evolutionarily Stable Community). In contrast, communities assembled by pure diversification (blue in Figure 6) tend to be characterized by low diversity and higher levels of generalism. These communities are often trapped at undersaturated local evolutionary states, and no matter the rate of (small) mutations, they cannot escape local optima without incoming dispersal of new trait values.

**Figure 6:**
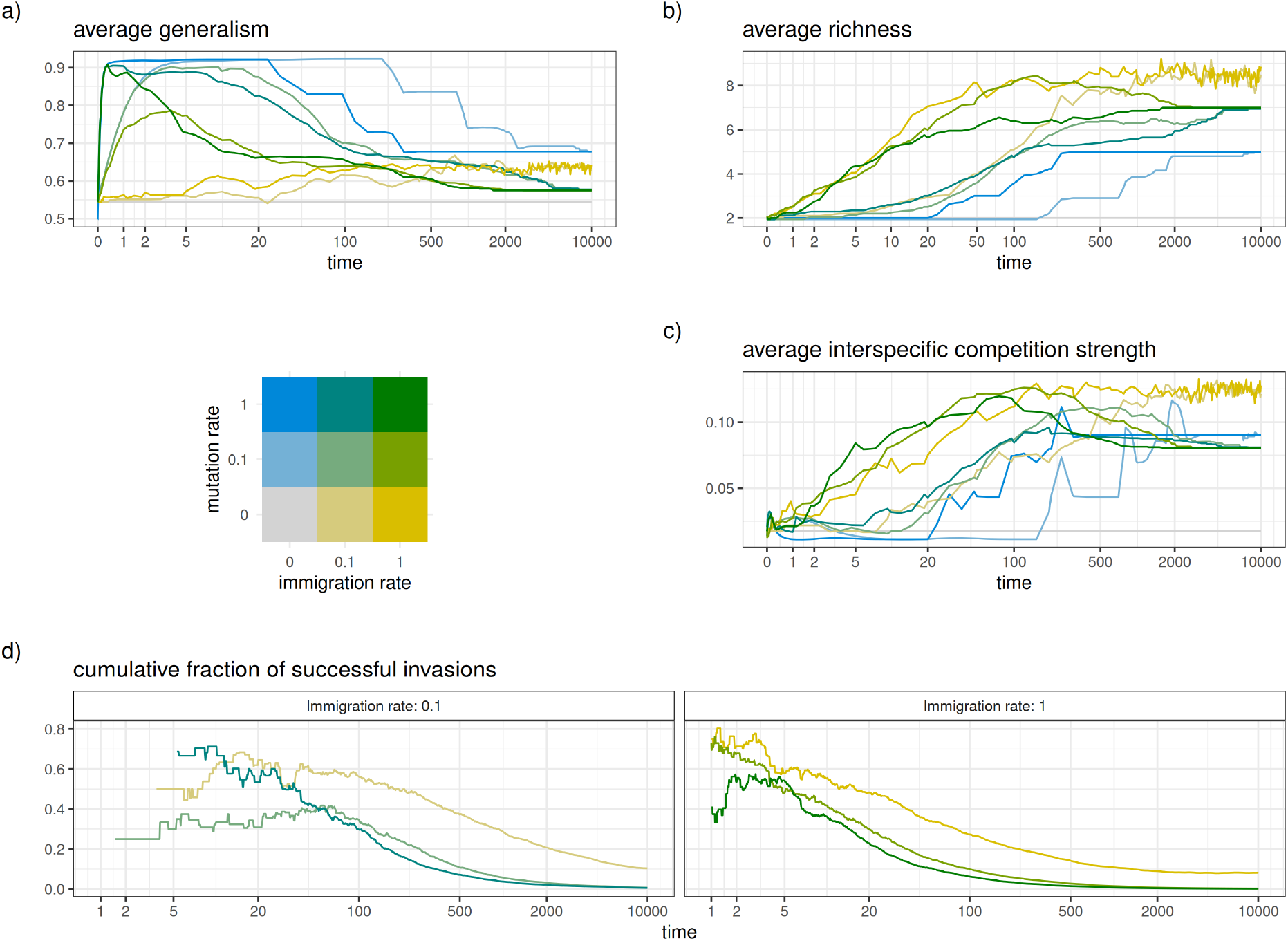
Evolution and immigration interact to jointly affect community assembly. Changes over time in: average generalism (a), average richness (b), and mean strength of competition (c) under different regimes of immigration and evolution. The bivariate color legend identifies the assembly regime: blue evolution and diversification alone, yellow no-evolution communities assembled through immigration only, and green - high rates of both evolution and immigration. Panel (d) shows the fraction of successful invasion events for three different evolution rates. Values are averages of 20 stochastic trajectories for each regime, time axes are pseudolog-transformed. Parameters not displayed on the plots: resource width = 6.5, trade-off *δ* = 2.0, *h* = 5*, m* = 0.2.

Finally, evolution rates affect the invasibility of communities that are subject to immigration (Figure 6, panel D). The fraction of successful immigration events decreases with time and richness, as the niches become increasingly occupied, but evolution accelerates this process (for a given rate of immigration *λ*). This occurs because adaptation of residents leaves fewer ecological opportunities for later-arriving migrants, a pattern known as community monopolization (De Meester et al., 2002). An increase in generalism does not appear to be the cause of this monopolization pattern. In fact, generalism tends to decline throughout the process, and even faster in evolutionary communities. Monopolization is not linked to higher richness either, since richness accumulates faster in immigration communities. Instead, monopolization appears to be due to evolution of niche positions *µ* whose divergence leads to better utilization of the resource axis: competition among morphs therefore declines in mixed assembly communities (green) compared to non-evolutionary ones (yellow) for a given immigration rate (Figure 6b).

## Discussion

We have investigated how evolution and immigration jointly affect the assembly and structure of consumer communities with varying degrees of generalism. Our findings highlight how resource availability, biological trade-offs, and modes of assembly constrain the evolution of generalism and modulate the tension between generalism and richness. First, the role of resource availability as an evolutionary driver of both diversification and greater generalism is consistent with empirical evidence that high-resource environments lead to both higher richness and broader niches (Costa-Pereira et al., 2019; Jones and Post, 2013). Given a large enough resource environment, we first observe evolution towards greater generalism of a single morph, then disruptive selection leading to diversification into several morphs. Increasingly larger resource axes favor diversity, as expected from classical fixed-width niche packing models (Case, 1981; MacArthur, 1969).

However, ecological trade-offs also play a role in constraining emerging richness and the degree of generalism. Generalism-efficiency trade-offs have occasionally been observed in nature for example in aphid parasitoids and hummingbirds Straub et al. (2011); Weinstein and Graham (2017) and several hypotheses have sought to explain the genetic basis of such costs to generalism: antagonistic pleiotropy, mutation-accumulation (Kawecki et al., 1997; Whitlock, 1996), and reduced evolvability (Bono et al., 2020). Despite a rich theoretical understanding of their central role in generalism evolution (Egas et al., 2004; Levins, 1962; Wickman et al., 2019), little empirical information exists on the actual shape of these trade-offs as they remain difficult to characterize (Kassen, 2002). Our exploration shows that the shape of fitness values across resources (parameter *δ*) also modulates the richness-generalism tension at the community level. When biology imposes a strong cost on generalism, diversification and specialism are favored over fewer, more generalist phenotypes (for the same resource environment, Fig 4a-b).

Contrary to discrete-resource models (Egas et al., 2004; Wickman et al., 2019), we observe the evolutionary emergence of a specialist-generalist pair as a possible outcome of evolutionary dynamics. Moreover, we identify multiple possible evolutionary endpoints for a same set of parameters, characterized by either fewer generalist types or numerous specialist types (Figure 4). This negative association between richness and generalism aligns with the negative correlation between intraspecific trait variation and diversity reported by (Barabás and D’Andrea, 2016; Barabás et al., 2022), and here we further demonstrate that it emerges even when both richness and generalism are products of the evolutionary process. It is also worth remarking that while generalism and intraspecific variation are related concepts, they are conceptually different and practically not always interchangeable. In fact, population niche width can be partitioned into a within-phenotype component (generalism) and a between-phenotype component (individual variation, Futuyma and Moreno (1988)). Whilst zero variation simply implies that all individuals exhibit the mean trait, there is a constraint on minimum viable generalism: when niche widths become very small, resource acquisition eventually becomes insufficient (Figure 1C). A corollary is that, even when niche width is allowed to vary, there is an upper limit to feasible ecological richness for a given resource environment (Germain et al., 2025). The niche variation hypothesis posits that generalism and intraspecific trait variation are correlated in natural populations, but evidence supporting it is mixed (Bolnick et al., 2007; Meiri et al., 2005).

Alternative evolutionary stable states at the community level have been observed in several models of eco-evolution (Ferriere and Legendre, 2013; Kremer and Klausmeier, 2017; Yacine et al., 2021), suggesting that founder effects, immigration events and stochasticity in mutations can crucially make evolutionary outcomes contingent. This is compatible with experimental and observational evidence illustrating how diversification is sensitive to immigration and can lead to different communities depending on the invaders’ order of arrival (Fukami et al., 2007; Gillespie, 2004). We show how, no matter the rate of evolution, purely evolutionary communities could not escape evolutionary dead-ends where increased generalism prevents further diversification a finding strikingly consistent with experimental evidence demonstrating that niche generalism reduces propensity for adaptive radiation in *Pseudomonas* (Flohr et al., 2013). Whilst these evolutionary local optima are a well-known concept at the level of genes and traits (Bank, 2022), their importance at higher levels of organization remains under-appreciated.

At the other end of the assembly spectrum, pure immigration communities manage to be both more generalist and richer than the global Evolutionarily Stable Community, thus occupying more fully the available resource axis and competing more strongly (Figure 6, panels A-C). We attribute this pattern to specialist consumers being more successful in invading the community by occupying small available niches, unlike the food web model of (Morris et al., 2021) where immigration of diverse preys favored more generalist consumers. Moreover, immigration communities in Figure 6 satisfy ecological conditions of coexistence but are not evolutionary endpoints, and multiple evolutionary models show that diversity at evolutionary stable points is generally lower than maximum ecological packing (Edwards et al., 2018; Rubin et al., 2023; Shoresh et al., 2008). Our results show that this pattern is conserved when generalism is an additional evolutionary variable, and that it is actually reinforced by the existence of non-global ESCs. We show that the influx of migrants must be sufficiently high relative to the mutation rate to transiently maintain the community at a richness level above that of the Evolutionarily Stable Community. This observation is ecologically relevant since more diverse communities are likely to consume available resources more efficiently (MacArthur, 1969) and, more broadly, to provide more ecosystem services (Cardinale, 2011; Hector and Bagchi, 2007; Maestre et al., 2012) but less rich communities may exhibit more robust coexistence (Lepori et al., 2024).

Finally, mixed assembly scenarios (fast evolution and high dispersal) are the most rapidly converging to globally uninvasible Evolutionarily Stable Communities (Edwards et al., 2018), aligning with the notion that immigration can help bring the adapted characters necessary to more rapidly fill available niches in an ecosystem (Richards et al., 2021). These mixed assembly regimes also display a typical pattern of niche monopolization where early evolution prevents successive invasion (De Meester et al., 2002), in line with empirical evidence from diversifying organisms at different scales demonstrating how the filling of available niches hinders both further diversification and immigration success (Fukami et al., 2007; Gillespie, 2004; Pantel et al., 2015). Our results suggest that the evolutionary divergence of niche positions, rather than the evolution of greater generalism, appears to be the main driver of community monopolization.

### Concluding remarks

Leveraging a relatively unexplored approach similar to the model of Ackermann and Doebeli (2004), which marries a continuous resource axis with varying niche widths, our study bridges ideas from different modeling traditions to provide a more unified understanding of how evolution and dispersal shape generalist-specialist communities. We validate findings from eco-evolutionary community assembly frameworks (such as resource-richness relationships, patterns of ecological versus evolutionary richness and community monopolization) and extend them through the explicit incorporation of varying degrees of generalism. Conversely, classical notions from studies of generalism such as the central role of allocation trade-offs are here generalized to species-rich communities and embedded within a community assembly context.

We further highlight the importance of alternative evolutionary states and local attractors of community structure, some of which are only reachable through immigration an under-explored facet of the interaction between local adaptation and regional dispersal dynamics. Whilst multistability is the object of much study in community ecology, where a large body of literature exists on priority effects and tipping points (Aguadé-Gorgorió et al., 2024; Stroud et al., 2024), these concepts deserve greater attention in the field of evolution (Dakos et al., 2019; Nosil et al., 2021). Given the accumulation of examples of eco-evolutionary dynamics, another important point is whether or not evolution maintains less diversity compared to communities based on ecological filtering, as several theoretical results seem to suggest (Edwards et al., 2018; Rubin et al., 2023) an idea somewhat in contrast with the concept of evolutionary rescue over shorter time-scales (Bell and Gonzalez, 2009; Gomulkiewicz and Holt, 1995). Our study, which explores assembly along a continuum of regimes, supports this prediction while also cautioning against the simplistic dichotomy of evolutionary and non-evolutionary communities. In fact, one can argue that the dynamics of assembly warrant equal attention as the final states: the fact that some ESCs might only be attainable under specific regimes might limit their practical relevance. These considerations highlight the need for frameworks that embrace an integrative perspective on community assembly across spatial, ecological, and evolutionary dimensions (Leibold et al., 2022a; Loeuille, 2019).

Finally, these findings open up exciting future directions for the study of the evolution of generalism in species-rich communities. As far as theory is concerned, one could envision an extension to this model in a full metacommunity framework that considers patches with distinct optima (Norberg et al., 2012). From an empirical standpoint, our results point to the value of community assembly experiments that integrate local adaptation, diversification, and immigration in suitable settings (e.g., the microbial systems in Flohr et al. (2013); Rainey and Travisano (1998)), which could test predictions of how assembly regimes shape patterns of community richness and generalism.

## Statement of Authorship

V.J.L.: conceptualization, methods development, model analysis, coding simulation, data visualization, writing (original draft), writing (review and editing); N.L.: conceptualization, methods development, supervision, writing (review and editing); R.P.R.: conceptualization, methods development, funding acquisition, supervision, writing (review and editing). This work has been funded by the Swiss National Science Foundation grant 31003A 182386 awarded to R.P.R..

## Data and Code Availability

All code (Julia simulation code and R plotting scripts) and outputs needed to reproduce the results and figures are available on Zenodo (https://doi.org/10.5281/zenodo.23974353).

