## Supplementary Materials for "Interplay of immigration and coevolution constrains the structure of specialist-generalist communities"

[1]Department of Biology – Ecology and Evolution, University of Fribourg

Chemin du Musée 10, CH-1700 Fribourg, Switzerland.

Institute of Ecology and Environmental Sciences, IEES

F-75005 Paris, France.

#### S1: From an explicit resource-consumer model to an implicit Lotka-Volterra model

Consider the classical MacArthur resource-consumer model for a logistically regenerating resource on a continuous axis $R(x)$ that is consumed by a discrete set of consumers $N_1, ..., N_i..., N_S$, with consumption niches $f(x|\mu_i\sigma_i)$. The region $R$ denotes the interval where resources are present, which depends on the choice of $\phi(x)$. Other symbols are described in the main text (Methods section and Box 1).

$$\frac{\mathrm{d}R(x)}{\mathrm{d}t} = R(x)\left(\phi(x) - \psi R(x) - \sum_i f(x|\mu_i, \sigma_i) \cdot N_i\right) \tag{1}$$

$$\frac{\mathrm{d}N_i}{\mathrm{d}t} = N_i\left(-m_i + \int_R R(x) \cdot f(x|\mu_i, \sigma_i)\ dx\right) \tag{2}$$

Where

$$\phi(x) = \begin{cases} h\ , & -l < x < +l \\ 0\ , & \text{otherwise} \end{cases} \tag{3}$$

$$f(x|\mu_i, \sigma_i) = a_i\ \mathrm{Exp}\left(-\frac{(x-\mu_i)^2}{2\sigma_i^2}\right) \tag{4}$$

It is possible to translate this model into an implicit model for consumer competition (MacArthur, 1972). To this end, we calculate the resource levels at any point on the axis for a given consumer community by assuming that resources always quickly reach equilibrium (separation of timescales assumptions). We here use $f_i(x) := f(x|\mu_i, \sigma_i)$ to simplify notation

$$R^*(x) = \frac{1}{\psi}\left(\phi(x) - \sum_i f_i(x) \cdot N_i\right) \tag{5}$$

If we further assume that the resource self-limitation parameter is $\psi = 1$, we can now substitute

$R^*$ into the consumer equation (2):

$$\frac{1}{N_i}\frac{\mathrm{d}N_i}{\mathrm{d}t} = -m_i + \int_R \phi(x) f_i(x)\ dx - \sum_j \int_R f_i(x) f_j(x)\ dx \cdot N_j \tag{6}$$

and we recover the well-known Lotka-Volterra dynamics by defining $r_i := -m_i + \int \phi(x) f_i(x) dx$ and $\alpha_{ij} := \int f_i(x) f_j(x) dx$ We now have a Lotka-Volterra model for competition between the consumers where resources are only implicit.

$$\frac{1}{N_i}\frac{\mathrm{d}N_i}{\mathrm{d}t} = r_i - \sum_j \alpha_{ij} N_j\ , \quad j = 1, .., i, .., S \tag{7}$$

#### S2: Checking the assumption of resource non-extinction using a discretized version of the resource-explicit model

The substitution operated in appendix S1 works under the assumption that the resource level $R^*$ is non-negative along the entire axis for the existing consumer community. However, when consumer pressure is high (for example when consumer mortality $m$ is low) some of the corresponding resources may implicitly become extinct (and even negative, which is biologically not meaningful). When $R^* < 0$, the implicit Lotka-Volterra ceases to be a perfect equivalent of the dynamics in the explicit resource-consumer model.

To test whether this would affect our conclusions, we have analyzed a discrete-resource equivalent formulation of the full model (Case and Casten, 1979). By discretizing the continuous resource axis R(x) into to a set of $K$ resources $R_1, R_2, ..., R_k$, where $R_j = R(x_j)$ and their spacing is $\Delta x$, while keeping consumer traits continuous, one then obtains the following model for $K$ resources and $S$ consumers:

$$\frac{dR_j}{dt} = R_j \left( \phi(x_j) - \psi R_j - \sum_{i=1}^{S} f(x_j \mid \mu_i, \sigma_i) \cdot N_i \right) \tag{8}$$

$$\frac{dN_i}{dt} = N_i \left( -m + \sum_{j=1}^{K} R_j \cdot f(x_j \mid \mu_i, \sigma_i) \cdot \Delta x \right) \tag{9}$$

The advantage here is that this discretized system can be written in generalized Lotka-Volterra form for both resources and consumers $dX/dt = X \cdot (\rho - AX)$, where $X = [R_1, ..., R_K, N_1, ..., N_S]$, and the interaction matrix $A$ is a block matrix describing interactions both within and between the resource and consumers. Notice how the consumer-consumer block of $A$ is empty: in this explicit model, competition only happens via the shared resources:

$$A = \begin{pmatrix} \psi I_K & F \\ -\Delta x F^\top & 0 \end{pmatrix}, \quad F_{ji} = f(x_j \mid \mu_i, \sigma_i), \tag{10}$$

Similarly the "growth" vector of this system contains positive resource regeneration rates and negative consumers background mortality (as consumers die in the absence of resources):

$$\rho = (\phi(x_1), \ldots, \phi(x_K), -m, \ldots, -m). \tag{11}$$

For a rare mutant consumer with trait $\mu_m, \sigma_m$, the invasion fitness in this discrete model depends on its own mutant traits, and on the resident trait set via the resource abundances $R^*(\boldsymbol{\mu_r}, \boldsymbol{\sigma_r})$

$$w_m = -m + \sum_{j=1}^{K} R_j^* \cdot f(x_j|\mu_m, \sigma_m)\Delta x. \tag{12}$$

This model is a direct discrete equivalent of the model in Appendix S1, and produces results perfectly consistent with the implicit Lotka-Volterra model presented in the paper, when the ecological equilibrium for the entire system $X^*$ is computed by solving the linear system $X^* = A^{-1}\rho$ (which allows negative resource levels $R_i < 0$). Conveniently, one can now also set up the ecological equilibrium resolution as a Linear Complementarity Problem (Takeuchi, 1996), a class of linear optimization problems which have algorithmic solutions allowing us to find a non-negative equilibrium satisfying $X^* \geq 0$ for both consumers and resources (i.e., extinct resources are exactly 0 and the non-zero abundances are computed correctly). This allows for studying the ecological and evolutionary dynamics while ensuring biologically consistent model behavior even when some of the resources are entirely drained. The code to run this full explicit model is included in the software folder.

Exploring this discrete explicit model indeed confirmed that the implicit Lotka-Volterra ceases to be a perfect approximation of the full model when some of the resources become extinct. Yet, we observe qualitatively similar patterns when using the LCP method to guarantee meaningful resource values when extinct. Namely, we find qualitatively similar effects of the resource width and tradeoff value on the position and stability of the evolutionary strategy. Moreover, we confirm the appearance of alternative evolutionary stable states, emergence of a generalist-specialist pair, and the existence of locally stable, non-globally uninvadable Evolutionary Stable Communities.

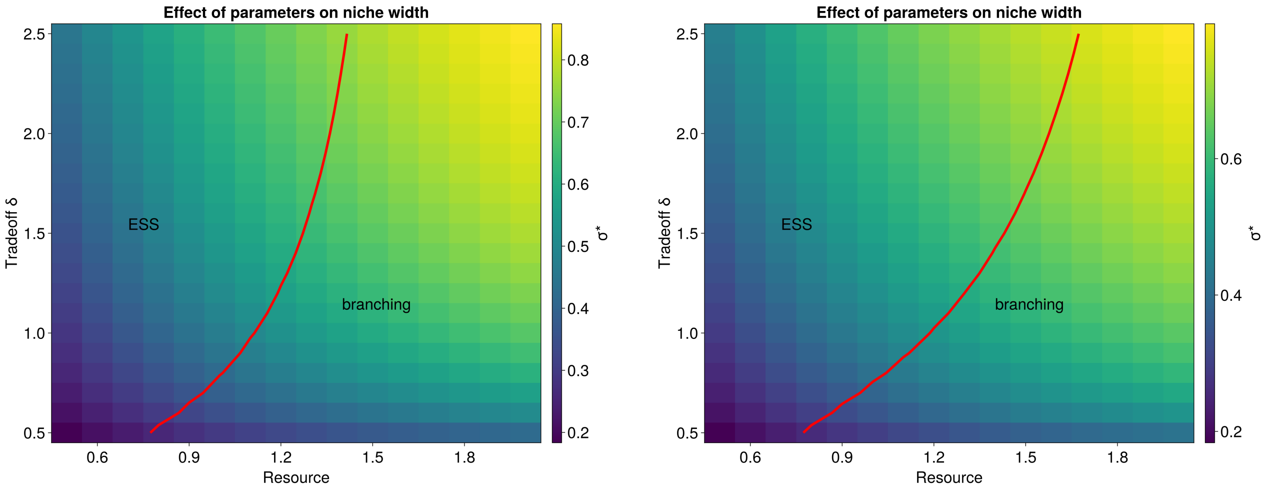


(a) (b)

Figure 1: *Location and stability of the monomorphic evolutionary equilbrium. Both panel are produced with the explicit, discrete resource model (here* $K = 40$*). The left panel uses this full model but resolving* $X^* = A^{-1}\rho$*. This model yields virtually perfectly equivalent results to the implicit Lotka-Volterra model with continuous resources (cf. left panel with Figure 2 in the main text). The right figure is also produced with the explicit discrete model but using the LCP method to compute biologically correct, non-negative resource abundances. Compared to the results obtained in the main text using the implicit model, the singular strategy remains stable (no branching) for slightly larger resource domains (for a given value of* $\delta$*). Despite this shift, the qualitative effects of the resource axis and tradeoff shape remain comparable.*

However, a complete exploration of this full model is made difficult by computational efficiency and numerical stability reasons: mainly, the LCP method often fails to solve the equilibrium for large systems with close values, where the domain of coexistence is small - such as in the regions near branching points. Yet, these results are reassuring as they suggest that the implicit Lotka-Volterra model is a reasonable approximation even under those scenarios (though not a perfect equivalent). Most importantly, the main patterns we observe in the main text are not simply a consequence of (biologically unrealistic) negative resource levels.

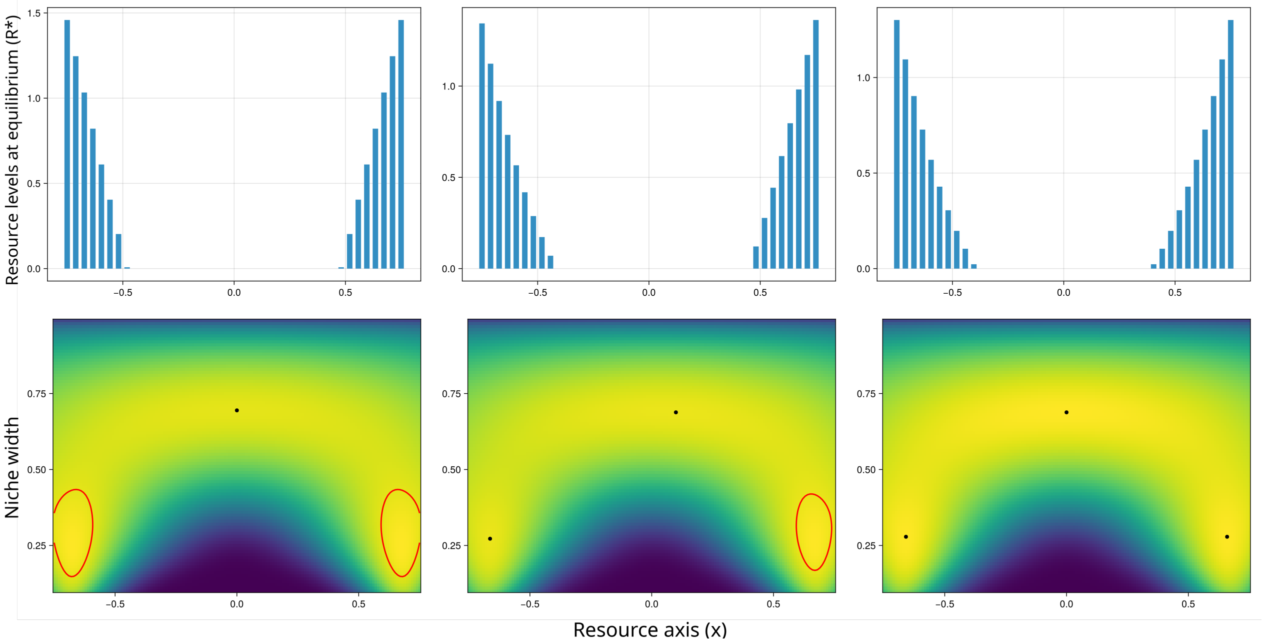


Figure 2: *Three alternative evolutionary stable states. The explicit, discrete-resources model with the LCP method similarly produces several alternative evolutionary endpoints for a same set of parameters, including specialist-generalist coexistence. Here we show 3 possible endpoints of evolution* $\mu^*, \sigma^*$ *(black dots, richness* $S^* = 1, 2, 3$*) for a same combination of parameters. Only the 3-morph coalition is global Evolutionarily Stable Community - red circles represent zero-contour lines highlighting small areas of the fitness landscape where* $w_m > 0$*. For each configuration, we show the ecological equilibrium levels for the discrete resources* $R_j^*$ *(here, we chose* $K = 40$ *for a reasonable approximation of a continuous resource axis case). Notice how some of them are entirely consumed but importantly, never negative. Moreover, we observe that the the consumer richness observed is comparable to that found in the implicit, continuous resource model. Parameters for this Figure are* $\delta = 2$*, resource width* $= 1.5$*.*

#### S3: Properties of the monomorphic singular strategy

We here show that the singular strategy for the niche position can be determined analytically. Recall the following definitions from the Lotka-Volterra model, in the case of a boxcar-like resource regeneration function $\phi(x)$ as in the main text:

$$f(x,\mu,\sigma) = \left(1-\sigma^{\delta}\right)^{1/\delta} \exp\left(-\frac{(x-\mu)^2}{2\sigma^2}\right) \tag{13}$$

$$\alpha(\mu_1,\mu_2,\sigma_1,\sigma_2) = \int_{-l}^{l} f(x,\mu_1,\sigma_1)\, f(x,\mu_2,\sigma_2)\, dx \tag{14}$$

$$\phi(x) = \begin{cases} h, & |x| \leq l \\ 0, & |x| > l \end{cases} \tag{15}$$

$$r(\mu_1,\sigma_1) = -m + \int_{-l}^{l} f(x,\mu_1,\sigma_1)\, \phi(x)\, dx \tag{16}$$

After integrating we get these forms for the competition coefficients and the consumers' growth rates - note the appearance of erf(x) functions due to integrating gaussians on a closed domain.

$$\begin{aligned} \alpha = {} & \frac{\sqrt{\pi/2}\,\sigma_1\sigma_2\left(1-\sigma_1^{\delta}\right)^{1/\delta}\left(1-\sigma_2^{\delta}\right)^{1/\delta}}{\sqrt{\sigma_1^2+\sigma_2^2}} \exp\left(-\frac{(\mu_1-\mu_2)^2}{2(\sigma_1^2+\sigma_2^2)}\right) \\ & \times \frac{-\mu_2\sigma_1^2-\mu_1\sigma_2^2+l(\sigma_1^2+\sigma_2^2)}{\sqrt{\left(\mu_2\sigma_1^2+\mu_1\sigma_2^2-l(\sigma_1^2+\sigma_2^2)\right)^2}} \operatorname{erf}\left(\frac{\sqrt{\left(\mu_2\sigma_1^2+\mu_1\sigma_2^2-l(\sigma_1^2+\sigma_2^2)\right)^2}}{\sqrt{2}\,\sigma_1\sigma_2\sqrt{\sigma_1^2+\sigma_2^2}}\right) \\ & + \frac{\mu_2\sigma_1^2+\mu_1\sigma_2^2+l(\sigma_1^2+\sigma_2^2)}{\sqrt{\left(\mu_2\sigma_1^2+\mu_1\sigma_2^2+l(\sigma_1^2+\sigma_2^2)\right)^2}} \operatorname{erf}\left(\frac{\sqrt{\left(\mu_2\sigma_1^2+\mu_1\sigma_2^2+l(\sigma_1^2+\sigma_2^2)\right)^2}}{\sqrt{2}\,\sigma_1\sigma_2\sqrt{\sigma_1^2+\sigma_2^2}}\right) \end{aligned} \tag{17}$$

and similarly for the consumers growth rates

$$r = -m + h\sqrt{\frac{\pi}{2}}\sigma_1 \left(1 - \sigma_1^\delta\right)^{1/\delta} \left[\mathrm{erf}\left(\frac{l - \mu_1}{\sqrt{2}\sigma_1}\right) + \mathrm{erf}\left(\frac{l + \mu_1}{\sqrt{2}\sigma_1}\right)\right] \tag{18}$$

We will then plug these expressions into the following expression for the fitness of a rare mutant for a scenario with only one resident, where we let 1 denote the mutant and 2 denote the resident:

$$\omega(\mu_1, \sigma_1|\mu_2, \sigma_2) = r(\mu_1, \sigma_1) - \alpha(\mu_1, \mu_2, \sigma_1, \sigma_2) \cdot N_2^* \tag{19}$$

where $N_2^* = r(\mu_2, \sigma_2)/\alpha(\mu_2, \mu_2, \sigma_2, \sigma_2)$. We now consider evolutionary gradient in the niche position, calculated as the derivative of equation (19) with respect to the mutant position $\mu_1$, evaluated at point $\mu = \mu_1 = \mu_2$, $\sigma = \sigma_1 = \sigma_2$. With help from the computer algebra software Mathematica, taking the closed forms (17,18) and derivating yields the following expression 20, for which we then need to find the root, i.e. the point $\mu^*$ where the evolutionary gradient vanishes.

$$\left.\frac{\partial \omega}{\partial \mu_1}\right|_{\substack{\mu_1=\mu_2 \\ \sigma_1=\sigma_2}} = \left(e^{2l\mu/\sigma^2} - 1\right) \cdot \frac{1}{2\sqrt{\pi}\,\sigma\left[\mathrm{erf}\left(\frac{l-\mu}{\sigma}\right) + \mathrm{erf}\left(\frac{l+\mu}{\sigma}\right)\right]} \Bigg[ - 2A\, e^{-(l+\mu)^2/(2\sigma^2)} \left(\mathrm{erf}\left(\tfrac{l-\mu}{\sigma}\right) + \mathrm{erf}\left(\tfrac{l+\mu}{\sigma}\right)\right) + e^{-(l+\mu)^2/\sigma^2} \left(1 + e^{2l\mu/\sigma^2}\right) \left(\sqrt{2}\,A\left(\mathrm{erf}\left(\tfrac{l-\mu}{\sqrt{2}\,\sigma}\right) + \mathrm{erf}\left(\tfrac{l+\mu}{\sqrt{2}\,\sigma}\right)\right) - 2m\right)\Bigg] = 0 \tag{20}$$

where $A = h\sqrt{\pi}\,\sigma_2\,(1 - \sigma_2^\delta)^{1/\delta}$. We can see that there is an obvious solution at point $\mu^* = 0$ which makes the first term zero, and Mathematica confirms that this is the only solution $\mu^* \in \mathbb{R}$ (there is a second, imaginary solution). This is expected given that the resource is fully symmetric, and confirms our numerical findings (Figure 2 in the main text). Conversely, we were not able to find a closed form for $\sigma^*$ using the same procedure.

### S4: Effects of evolutionary rates on polymorphic ESC

Here we test how relaxing the assumption of similar rates of evolution $M$ for $\mu$ and $\sigma$ affects community dynamics. To this end, we rerun the simulations with $M_\mu = 0.1 \cdot M_\sigma$, and vice-versa. This does appear to affect the dynamics but not the endpoints reached under evolutionary assembly. Figure 3 shows that, while monomorphic evolutionary trajectories are altered by a change in the relative rates, the singular strategy towards which evolution converges remains the same (this is expected given that $M$ is a positive constant not affecting the root). Importantly, even at higher levels of richness, the evolutionary endpoints and their relative frequencies do not appear to differ substantially when either niche positions or niche widths evolve faster (Figure 4).

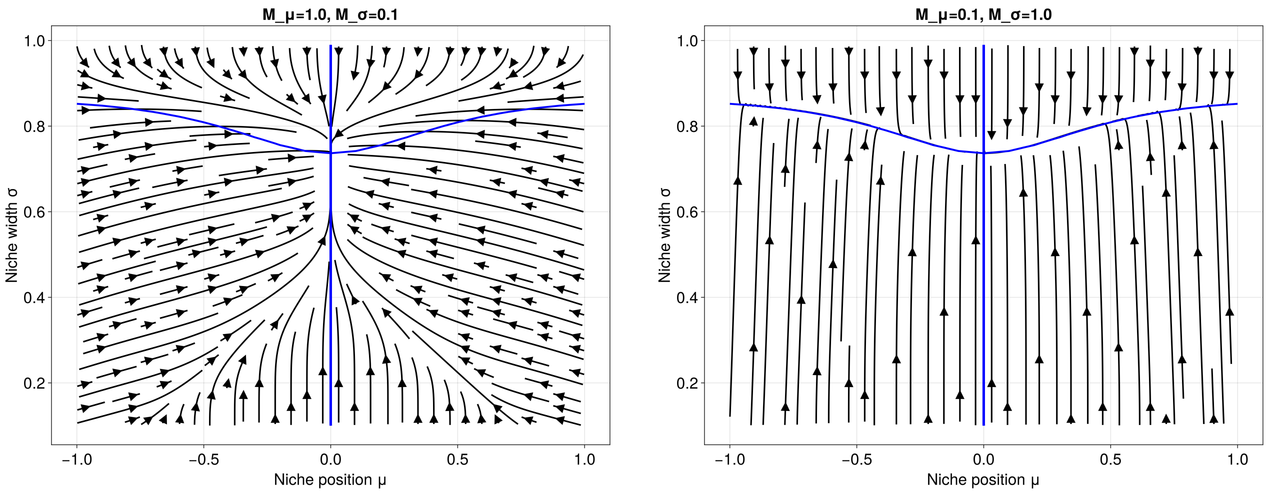


(a) (b)

Figure 3: The two diagrams show the evolutionary endpoints obtained for different rates of $M_\mu/M_\sigma$. Other parameters are kept the same as for Figure 2 in the main text.

Similarly, in the mixed-assembly scenarios (Figures 5 and 6), the communities emerging under different regimes over very long time horizons appear preserved despite imbalances in the evolutionary rates of $\mu$ and $\sigma$. Over long transient periods, however, we observe some differences between the dynamics: all else being equal, when $M_\sigma > M_\mu$, phases of high generalism in evolutionary and mixed communities persist for longer, and evolutionary communities may remain in low richness states for longer.

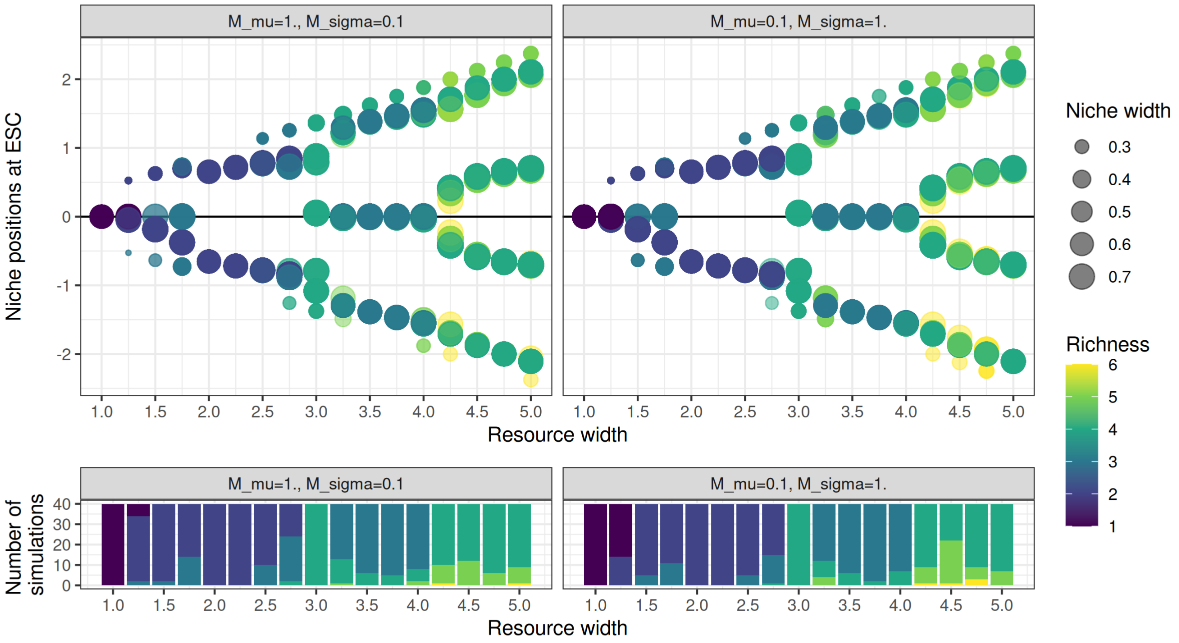


Figure 4: The two panels show the evolutionary endpoints obtained for different ratios of $M_\mu/M_\sigma$. Other parameters are kept the same as for Figure 5 in the main text.

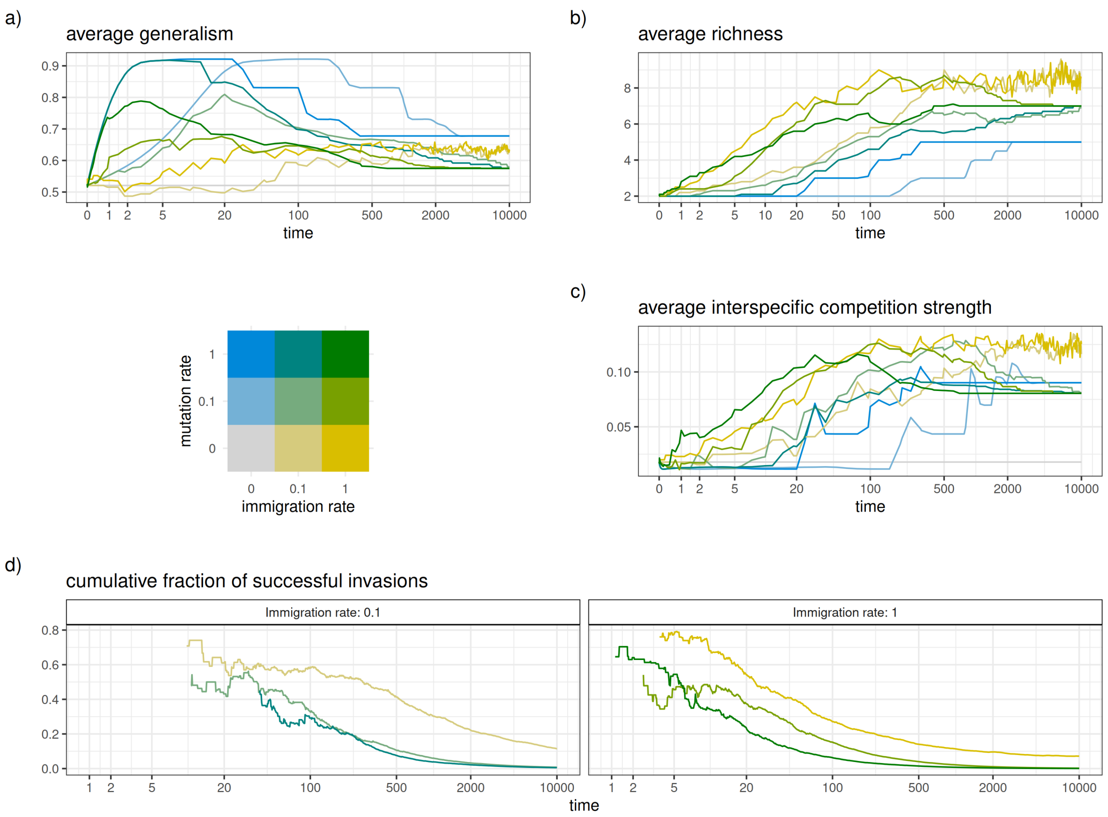


Figure 5: **Mixed assembly when niche positions evolve faster than niche widths**. Here, mutation rates in the color legend indicate $M_\mu$, with $M_\sigma = 0.1 \cdot M_\mu$. Other parameters are kept the same as for Figure 6 in the main text.

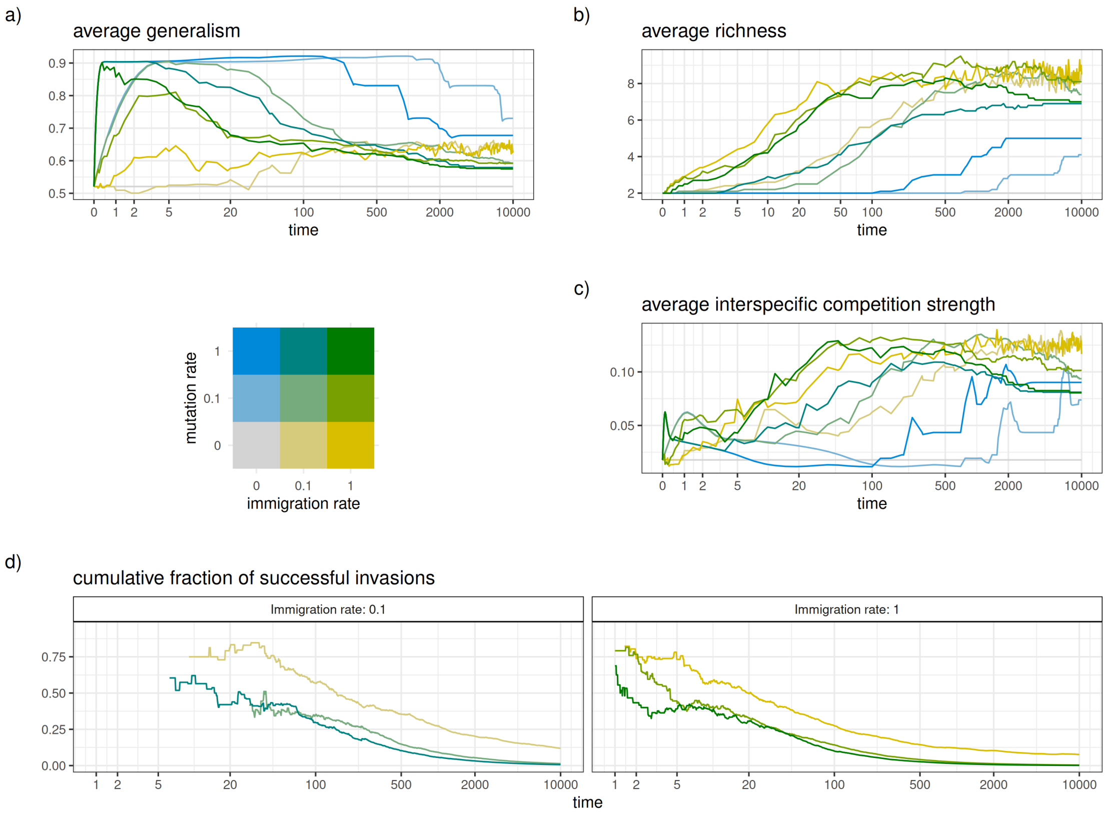


Figure 6: **Mixed assembly when niche widths evolve faster than niche positions**. Here, mutation rates in the color legend indicate $M_\sigma$, with $M_\mu = 0.1 \cdot M_\sigma$. Other parameters are kept as in Figure 6 in the main text.
